# Field evaluation of DNA amplification of human filarial and malaria parasites using mosquito excreta/feces

**DOI:** 10.1101/2019.12.23.880740

**Authors:** Corrado Minetti, Nils Pilotte, Michael Zulch, Tiago Canelas, Edward J. Tettevi, Francis B. D. Veriegh, Mike Yaw Osei-Atweneboana, Steven A. Williams, Lisa J. Reimer

## Abstract

**Background:** We recently developed a superhydrophobic cone-based method for the collection of mosquito excreta/feces (E/F) for the molecular xenomonitoring of vector-borne parasites showing higher throughput compared to the traditional approach. To test its field applicability, we used this platform to detect the presence of filarial and malaria parasites in two villages of Ghana and compared results to those for detection in mosquito carcasses and human blood.

**Methodology and principal findings:** We compared the molecular detection of three parasites (*Wuchereria bancrofti, Plasmodium falciparum* and *Mansonella perstans*) in mosquito E/F, mosquito carcasses and human blood collected from the same households in two villages in the Savannah Region of the country. We successfully detected the parasite DNA in mosquito E/F from indoor resting mosquitoes, including *W. bancrofti* which had a very low community prevalence (2.5-3.8%). Detection in the E/F samples was concordant with detection in insect whole carcasses and people, and laboratory tests showed that the risk of mosquito carcass cross-contamination with positive excreta when insects are held together in the device is minimal.

**Conclusions:** Our approach to collect and test mosquito E/F successfully detected a variety of parasites at varying prevalence in the human population under field conditions, including a pathogen (*M. perstans*) which is not transmitted by mosquitoes. The method shows promise for further development and applicability for the early detection and surveillance of a variety of pathogens carried in human blood.

**Author summary:** Molecular xenomonitoring of parasites or viruses using mosquitoes as “flying syringes” is a promising and non-invasive tool for early detection and surveillance of various pathogens, particularly in low prevalence settings, but there is a need for cost-effective and higher throughput alternatives. We recently developed a novel approach based on the collection of mosquito excreta/feces (E/F) using a superhydrophobic cone that directs the sample into a tube at the bottom of the collection cup. We tested this method’s ability to detect the presence of lymphatic filariasis, malaria, and mansonellosis from households in two endemic rural communities of Ghana and compared it to the molecular detection from blood and mosquito carcass samples from corresponding households. The detection of parasite DNA in mosquito E/F was successful for all three pathogens, and it showed good concordance with the detection in the insects’ whole carcasses. Given the successful detection of *Mansonella perstans*, which is not transmitted by mosquitoes, our tool shows promise for use as an alternative method for the early and non-invasive detection and surveillance of various pathogens in human blood, including those not strictly mosquito-borne, in endemic settings.

## Introduction

The detection of nucleic acids (DNA or RNA) from human pathogens in insect vectors (molecular xenomonitoring, MX) to assess their presence in a community is an important and promising approach to disease surveillance. More broadly, by using hematophagous arthropods as environmental samplers or ‘flying syringes’, any pathogen circulating in the blood of vertebrate hosts that is taken up in a bloodmeal can be detected by polymerase chain reaction (PCR) in the insect carcass (xenosurveillance) [1–4]. Since pathogen DNA or RNA can remain detectable for several days in arthropod bloodmeals [1,5], screening for pathogens which are not normally transmitted by a particular insect species is possible [6]. Examples include the detection of *Plasmodium vivax* DNA in the abdomen of blood fed *Culex* mosquitoes collected in the households of people with malaria [7], or the use of anopheline mosquitoes to detect Dengue viruses [8]. Molecular xenomonitoring offers an alternative to the standard approaches for disease surveillance: it can provide a real-time representation of the pathogens (parasitic, bacterial or viral) circulating in a population, it does not rely on collecting samples directly from people, and it can make use of insect samples collected for other purposes.

These advantages become clear in the context of diseases such as lymphatic filariasis (LF), which are nearing elimination in many countries. According to the strategy of the Global Programme to Eliminate Lymphatic Filariasis (GPELF), interruption of transmission following mass drug administration (MDA) is assessed via transmission assessment surveys (TAS) where either adult worm antigens or host antibodies are detected in school children [9]. However, since the sensitivity of parasite antigen detecting tests is reduced in low prevalence settings [10], the long term cost-effectiveness of this strategy and its ability to detect ongoing transmission has been put into question and an alternative approach may be needed. Molecular xenomonitoring has proven useful in the assessment of LF transmission or validation of elimination in many countries including Papua New Guinea [11], Ghana [12], Togo [13], Egypt [14], India [15], Sri Lanka [16] and American Samoa [17].

To implement MX in LF elimination programmes on a global scale, both the protocols for collecting different species of mosquitoes in different settings and the molecular diagnostic tools used for detection must be standardized. Furthermore, the levels of infection in vectors allowing for sustained transmission (accounting for vector density and biting rates) need to be clearly defined [18,19]. Currently, the cost-effectiveness and potential for scaling up MX using insect carcasses is negatively affected by the need to collect and process large numbers of samples [19]. Further constraining efforts, mosquitoes can only be pooled for DNA extraction in groups of up to 25 insects to avoid a loss of sensitivity in parasite detection [20]. Since such limitations can be reasonably applied to the MX of any pathogen, particularly in low prevalence settings, there is a need for more cost-effective alternatives ensuring high sensitivity of detection and a higher throughput.

The discovery that the DNA of ingested parasites can also be detected in the excreta/feces (E/F) of mosquitoes [21], regardless of whether they are competent vector species or not, has offered a novel and promising approach to MX. Insect E/F provide a ‘cleaner’ sample, minimizing the mass of mosquito DNA which may affect the amplification of the target organisms. Additionally, MX of E/F substantially increases the throughput of mosquito processing while maintaining sensitivity: it has been shown to successfully amplify pathogen DNA in the E/F from one positive mosquito within an E/F pool from up to 500 negative mosquitoes [22]. The use of mosquito E/F for MX purposes has recently been demonstrated for a variety of arboviruses [23–25], showing that viral RNA can also be detected in the field [26]. A recent laboratory study has also shown that *Plasmodium falciparum* RNA can be detected concomitantly in the excreta and saliva of infected *Anopheles stephensi*, with good concordance between the two types of sample [27]. However, to our knowledge, E/F collections for the detection of eukaryotic pathogens have yet to be conducted in a field setting.

To ensure a more efficient E/F collection from field-caught mosquitoes outside a laboratory setting, we recently developed a paper conical insert sprayed with a superhydrophobic coating which concentrates the E/F produced from blood fed mosquitoes housed either individually or in groups [28]. The insert can fit into paper cups of different sizes, is re-usable following a brief wash, and the E/F can be collected either into a microcentrifuge tube or onto an absorbent sample collection card for preservation. In the laboratory we successfully detected parasite DNA in the E/F of mosquitoes fed a variety of parasites (*Brugia malayi, P. falciparum* and *Trypanosoma brucei brucei*) [28], and our preliminary field tests in Ghana showed that the device can accommodate blood fed mosquitoes collected indoors without causing mortality to the insects. Using the superhydrophobic cone (SHC), or a similarly sprayed surface, in a passive mosquito trap could potentially lead to a community-wide multi-pathogen, non-invasive infection surveillance approach with minimal labor or cost.

Here we present an evaluation of this promising novel approach to MX of *Wuchereria bancrofti, P. falciparum* and *Mansonella perstans* in two rural communities of Ghana. The infection detection efficiency in mosquito E/F was compared to that determined using traditional approaches including MX using insect carcasses and parasitological examination of human samples. Our primary aim was to determine the multi-disease potential of this approach by evaluating the detection of both mosquito-borne and non-mosquito borne parasites and pathogens. Our secondary aim was to use these data to infer the spatial distribution of parasite positivity in the different specimens to help inform decisions as to whether a community-wide or more focal mosquito sampling is required.

## Methods

### Study communities

We conducted the study in the communities of Sekyerekura and Dugli, located in the Bole District, in the Savannah Region of Ghana. We previously surveyed all the households (24 in Sekyerekura and 47 in Dugli) in March 2017 as part of a study on the status of elimination and determinants of LF persistence in Ghana [29]. At the time of the survey, estimated *W. bancrofti* microfilaria community prevalences were 7.4% and 5.5% for Sekyerekura and Dugli, respectively. An additional finding was the presence of *M. perstans*. This neglected filarial parasite is transmitted by biting midges of the genus *Culicoides*, and recent data have shown that it is endemic in Ghana [30]. Communities in this part of the country are also highly endemic for *P. falciparum* malaria [31] and they are located in a dry savannah environment, with a single rainy season beginning in May and ending in October. In both villages, the dominant mosquito vectors of both LF and malaria are *Anopheles gambiae* s.s, *An. arabiensis* and *An. funestus* [31,32]. We conducted this study in October 2017, at the end of the rainy season. As part of the national program for LF elimination, MDA with ivermectin and albendazole took place in the communities two to four weeks before the study.

### Household mapping and selection

We mapped all the households in both villages by collecting the GPS latitude and longitude coordinates with tablets equipped with the Open Data Kit (ODK) application, and by assigning each a unique serial number. In each village, we then randomly selected twelve households for parasitological testing using the random number generator function in Microsoft Excel.

### Parasitological testing in selected households

In the selected households, we invited all people 1 year of age and above to take part in the parasitological survey. The presence of *W. bancrofti* adult worm and *P. falciparum* antigens was determined in consenting participants (and assenting children) by finger-prick blood testing using the Filariasis Test Strip (FTS) (Alere Inc.,Waltham, USA) and the OptiMAL-IT (Diamed GmBH, Switzerland) immunochromatographic tests, respectively, as per the manufacturer’s instructions. In addition, approximately 25 µl of blood from each participant was blotted onto a Whatman filter paper disk, which was dried and stored in bags with silica gel beads for later DNA extraction and testing for parasite DNA presence. During the night (2200h to 0100h), 2 ml of venous blood was also collected from consenting FTS-positive participants to enable detection of *W. bancrofti* microfilariae using the acetic acid fixation and counting chamber method, and to estimate the number of larvae per millilitre of blood [33]. For all consenting participants age and sex were also recorded. All participants which tested positive for malaria antigen were provided artemisinin combination therapy (ACT) with artemether lumefantrine by a registered nurse, following the Ghana national guidelines [34].

### Mosquito and E/F collections

In all assenting households from both villages, between 0500h and 0800h, we collected indoor resting mosquitoes from ceiling, walls, furniture and hanging clothes using a battery powered aspirator [35]. A total of 26 (56 sleeping spaces) and 49 (116 sleeping spaces) households were sampled twice in Sekyerekura and Dugli, respectively. Additionally, gravid and host-seeking mosquitoes were also collected using a combination of *Anopheles* and Box gravid traps and BG-sentinel traps as described previously [36]. In total, 12 collection nights were performed per trap type. At the time of collection all live female mosquitoes were transferred into paper cups covered with netting and transported to a central location for processing. Here, according to the household of collection, mosquitoes were sorted into paper cups equipped with the SHC positioned above a 1.5 ml microcentrifuge tube and were provided access to sugar-soaked cotton wool as previously described [28]. Mosquitoes were held in the E/F collection cups in pools per household (with pool sizes ranging from 1 to 26 insects) for approximately 35 hours, after which they were processed. The tubes with the E/F were sealed, with three holes made in the lid, and then stored in plastic bags containing silica beads. Mosquitoes were morphologically identified to the species complex level [37] and then stored individually in 1.5 ml collection tubes with silica beads for later molecular processing. The SHCs were rinsed with distilled water between collections and re-used, since laboratory trials previously confirmed the absence of DNA contamination when performing this rinse procedure [28].

### Parasite molecular detection

We extracted the DNA from mosquito E/F in the 1.5 ml collection tubes using the QIAamp DNA Micro Kit (QIAGEN, Germantown, Maryland) following a modified version of the manufacturer’s protocol as described elsewhere [38]. To test the possibility of detecting the presence of *W. bancrofti* or *P. falciparum* developing or infective stage, mosquito carcasses were separated into two separate aliquots, heads and thoraces (H&T) and abdomens (Abd), which were then pooled in groups from up to five mosquitoes. Carcass DNA was extracted using the DNeasy 96 Blood & Tissue kit (QIAGEN, Manchester, UK) as per the manufacturer’s instructions, with the following modification: before incubation in ATL lysis buffer at 56°C for 3 hours, the tissues were disrupted in the same buffer using steel beads in a TissueLyser (QIAGEN) shaker (5 minutes at 30 1/S frequency). Dried blood spots were extracted following a previously published protocol [39]. The presence of parasite DNA in the extracted samples was determined by real-time quantitative PCR (qPCR). The presence of *W. bancrofti* and *P. falciparum* DNA was tested using two novel assays targeting repetitive regions in the parasites’ genome and described elsewhere [40]. The presence of *M. perstans* was tested using a previously undescribed assay targeting the parasite ribosomal RNA internal transcribed spacer (ITS) region (available at the following address: https://www.ncbi.nlm.nih.gov/nuccore/EU272184.1?report=fasta). The newly designed primer sequences were as follows: 5’-ATAGACACCAACGAATATCACCGTTA-3’ for the forward and 5’-TCATTAACGAGCTTCCAAACAAATACA-3’ for the reverse. A previously published probe (6-FAM-TTCACTTTTATTTAGCAACATGCA-TAMRA) [41] was used for detection. All qPCR assays were run in 25 µl total volumes, each including 500 nM of each primer, 200 nM of probe, 1X TaqPath™ ProAmp™ Master mix (Applied Biosystems) and 5 µl of template DNA, with the following cycling conditions: 50°C for 10 min, 95°C for 2 min and 45 cycles of 95°C for 15 seconds and 60°C for 1 min. All samples were initially tested in duplicate. If one replicate was negative and the other was positive, then the sample was retested. A sample was considered positive either if it allowed for amplification in both the initial reaction replicates or it was positive in at least one replicate reaction upon retesting. As described elsewhere [38], highly sensitive digital PCR using the QuantStudio 3D Digital PCR instrument was used to retest E/F samples that were inconsistently positive upon initial testing, given the very small residual volumes available for retesting from these samples and the potential to improve detection when very low concentrations of target are present. Due to the prohibitive cost of digital PCR, qPCR was instead used to retest inconsistently positive blood and mosquito carcasses samples, as DNA extraction from these sample types produces considerably greater sample volumes.

The study design and sample collection workflow are depicted in Figure 1.

**Figure 1.**
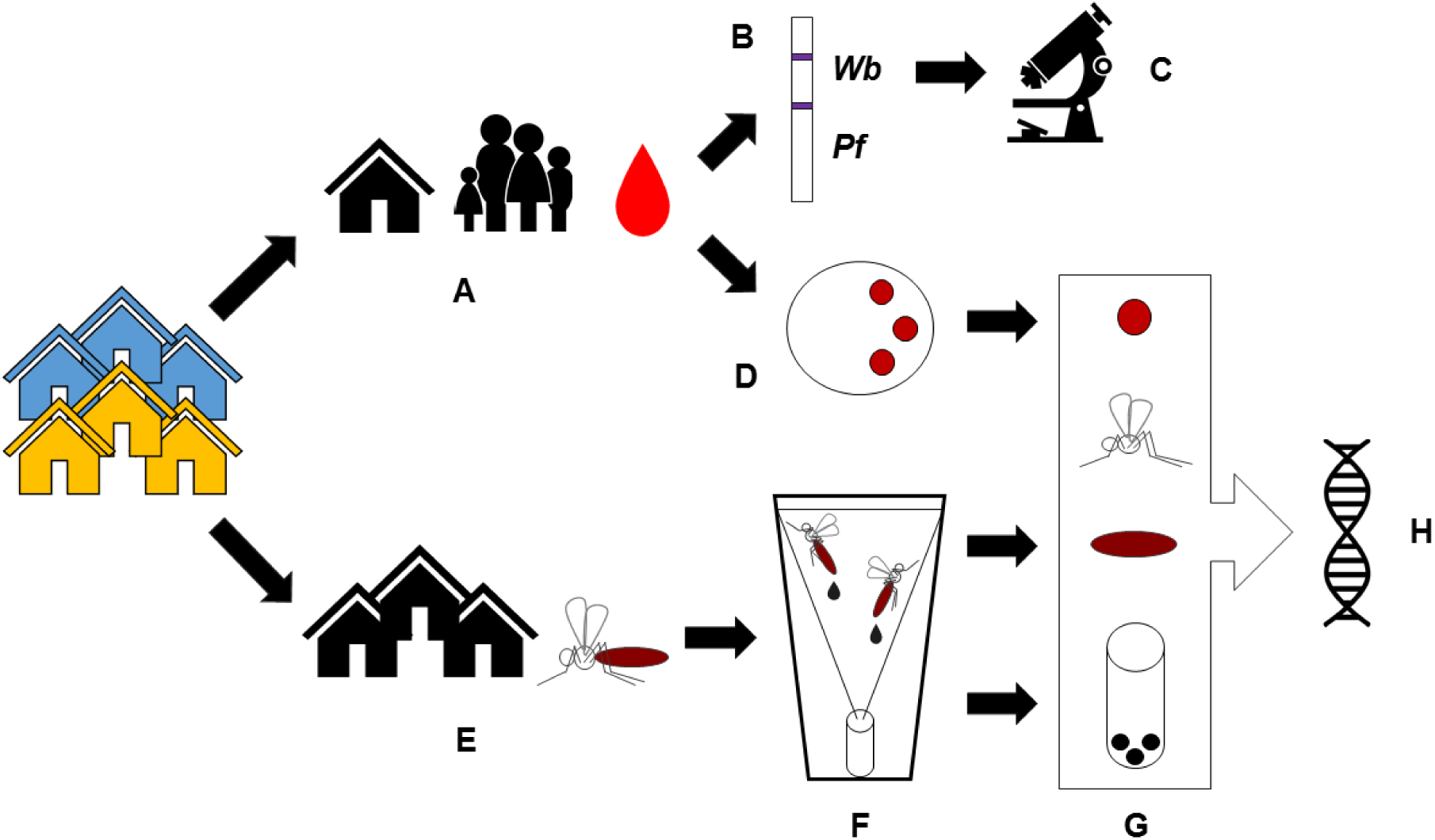
Study design and sample collection workflow. A) Twelve households were selected from each of the two communities, and blood was collected from all consenting individuals >1 y of age; B) The presence of filarial and malaria antigen was determined by immunochromatographic tests, and C) the presence of microfilaria was determined by microscopy in the night blood of people who tested positive for the filarial antigen; D) Blood spots were collected onto filter paper; E) Indoor resting female mosquitoes were collected from all households; F) Mosquitoes were held in collection cups fitted with the SHC to collect the E/F; G) Dried blood spots, mosquito E/F and mosquito carcasses were processed and H) DNA was extracted from the samples and the presence of *W. bancrofti, P. falciparum* and *M. perstans* determined by qPCR.

### Statistical analysis and mapping

Data were analysed using the IBM SPSS Statistics (version 22) and R software. Descriptive statistics were calculated for parasite infection prevalence in humans, and mosquito abundance and diversity. Parasite DNA prevalence in mosquito E/F and carcasses (overall and separate for H&T and Abd) was calculated using maximum likelihood estimates (with 95% confidence intervals) using PoolScreen 2.02 [42]. Associations in the DNA positivity between mosquito E/F and carcasses (overall and separate for H&T and Abd) and between mosquito E/F or carcasses and household members were calculated using the Pearson’s correlation coefficient r or the Chi-square test, respectively. Maps for household parasites’ positivity in human blood and mosquito carcasses and E/F were made in QGIS 3.x. Mosquito density was interpolated using the Inverse Distance Weighted method. In this method, sample points are weighted during interpolation such that the influence of one point relative to another declines with distance. We performed a spatial clustering analysis for each pathogen and mosquito density in both villages in the study site. Pathogen positivity was treated as a case-control point data, with cases being households positive for that pathogen and controls being households which were negative. Differences in K functions were calculated, and tolerance envelopes produced, then a global test of clustering using difference in K functions was applied by comparing cases and controls [43]. Clustering for mosquito density was assessed by using the Getis-Ord General G statistic. Using our dataset, the probability (with 95% confidence intervals) of finding a mosquito sample (either whole carcass or E/F sample) positive for *W. bancrofti* or *M. perstans* sampling 10 randomly selected houses was calculated using a hypergeometric distribution function.

### Ethical clearance

Ethical approval for the study was obtained by the Ethics Committee of the Liverpool School of Tropical Medicine, United Kingdom (<u>Research Protocol 17-035 A vector excreta surveillance system VESS to support the rapid detection of vector-borne diseases</u>) and the Council for Scientific and Industrial Research, Accra, Ghana. Members of the research team met with the district health officials and community leaders to explain the purpose of the study before enrolling the participants. Meetings were then held at the community level to explain the purpose of the study to all residents, and further details were explained to the selected household participants before they signed an informed consent form for themselves or on behalf of their children.

### Testing for mosquito carcass cross-contamination through E/F

To evaluate the possibility of cross-contamination of mosquito carcasses through exposure to E/F of other mosquitoes during the holding period in the SHC, we performed a series of laboratory tests. *An. gambiae* mosquitoes from the G3 colony (Liverpool School of Tropical Medicine) were fed cultured *P. falciparum* (strain 3D7) at 0.1% parasitemia. Five *P. falciparum*-exposed mosquitoes were then co-housed with 20 unexposed *A. aegypti* mosquitoes (Liverpool School of Tropical Medicine colony) in collection cups fitted with an SHC. This experiment was replicated five times. Co-housing occurred from 24 to 72 hours post exposure, at which point E/F samples were collected and mosquitoes were knocked down. DNA extraction and qPCR analysis for the presence of *P. falciparum* was then performed individually on mosquito carcasses as described in the sections above. Unexposed mosquitoes were similarly tested, however, extractions and qPCR testing occurred on whole insects. To verify the presence of *P. falciparum* signal within the potentially contaminating E/F, all E/F samples also underwent DNA extraction and qPCR testing as described in the sections above.

## Results

### Mosquito abundance and diversity

A total of 2,331 mosquitoes were collected across the two surveyed communities. The majority of collected mosquitoes (>95%) were anophelines, which included mostly *An. gambiae* sl (87%) followed by *An. funestus* sl (10.6%). The number and percentage of mosquitoes by species complex/genus and by collection method are reported in S1 and S2 Tables.

### Parasite prevalence in humans and in mosquito E/F and carcasses

The human prevalence and the DNA positivity in mosquito E/F and carcasses (overall and separate for H&T and Abd) of the three parasites in the two sampled communities are shown in Tables 1 to 3. None of the blood spots collected during the day tested positive for *W. bancrofti* DNA by qPCR but microfilaria was detected in night blood from three and two people in Sekyerekura and Dugli, respectively. The prevalence of malaria was high by both antigen (39-42.1%) and qPCR (72-82.4%) in both communities. No species other than *P. falciparum* was found in the samples. The filarial parasite *M. perstans* was also detected at a prevalence of 35.4-37.4% in participants’ blood by qPCR, and microfilaria were also confirmed in the night blood samples collected to screen for *W. bancrofti*.

**Table 1.** *W. bancrofti* prevalence in human blood, mosquito E/F and carcasses.

| Community | FTS %<br>(+/tested) | Mf %<sup>a</sup><br>(+/tested) and mean<br>density<sup>b</sup> | E/F DNA %<sup>c</sup><br>(+pools/tested) | Car DNA %<sup>c</sup><br>(+pools/tested) |
| --- | --- | --- | --- | --- |
| Sekyrekura | 17.7 (14/79) | 3.8 (3/14)<br>170 larvae/ml | 0.14 (0.0042-0.71)<br>(1/81) | 0.66 (0.19-1.55)<br>(5/197) |
| Dugli | 19.2 (15/78) | 2.6 (2/15)<br>40 larvae/ml | 0.20 (0.039-0.59)<br>(3/139) | 1.10 (0.61-1.80)<br>(17/383) |

FTS= filariasis test strip; Mf= microfilaria; E/F= excreta/feces; Car= whole carcass; <sup>a</sup>Community microfilaria prevalence, assuming those testing negative for antigen were also negative for mf, estimated as follows: (% FTS+ \* % mf+ in the FTS+); <sup>b</sup>Number of larvae per millilitre of blood; <sup>c</sup>Filarial DNA prevalence per mosquito (maximum likelihood and 95% CI) were estimated using PoolScreen2.

**Table 2.** *P. falciparum* prevalence in human blood, mosquito E/F and carcasses.

| Community | RDT %<br>(+/tested) | Blood qPCR %<br>(+/tested) | E/F DNA %<sup>a</sup><br>(+pools/tested) | Car DNA %<sup>a</sup> (+pools/tested) |
| --- | --- | --- | --- | --- |
| Sekyrekura | 45.1 (41/91) | 82.4 (75/91) | 0.42 (0.082-1.22)<br>(3/81) | 12.41 (9.62-15.63) (77/197) |
| Dugli | 39 (32/82) | 72 (59/82) | 0.67 (0.31-1.29)<br>(10/139) | 20 (17.38-22.93) (225/383) |

RDT=rapid diagnostic test; qPCR= real-time quantitative PCR; E/F= excreta/feces; Car= whole carcass; <sup>a</sup>Malaria DNA prevalence per mosquito (maximum likelihood and 95% CI) were estimated using PoolScreen2.

**Table 3.** *M. perstans* prevalence in human blood, mosquito E/F and carcasses.

| Community | Blood qPCR %<br>(+/tested) | E/F DNA %<sup>a</sup> (+pools/tested) | Car DNA %<sup>a</sup> (+pools/tested) |
| --- | --- | --- | --- |
| Sekyerekura | 37.4 (34/91) | 0.70 (0.21-1.65) (5/81) | 8.21 (6.03-1.08) (55/197) |
| Dugli | 35.4 (29/82) | 1.94 (1.21-2.91) (26/139) | 7.48 (6-9.17) (104/383) |

qPCR= real-time quantitative PCR; E/F= excreta/feces; Car= whole carcass; <sup>a</sup>Filarial DNA prevalence per mosquito (maximum likelihood and 95% CI) were estimated using PoolScreen2.

The DNA of the three parasites was successfully detected in mosquito E/F, with *M. perstans* found in the highest number of E/F pools (Table 1). Parasite DNA positivity in mosquito E/F was not associated with the number of mosquitoes in the pool. The prevalence of DNA positivity in mosquito carcasses was higher in the H&T aliquot than in the Abd aliquot for *W. bancrofti*, whereas the opposite was observed for *P. falciparum* and comparable positivity rates were observed between the two body parts for *M. perstans* DNA. Positive E/F for *W. bancrofti* and *P. falciparum* were only found from resting mosquitoes collected indoors, whereas *M. perstans* DNA was also detected in the E/F deposited by three mosquitoes collected using the CDC Box trap and one collected using the *Anopheles* gravid trap.

Parasite DNA positivity in mosquito E/F was significantly and positively associated with carcass positivity for *W. bancrofti* (Pearson’s r 0.119, *p*<0.01) and *M. perstans* (Pearson’s r 0.135, *p*<0.01), but not for *P. falciparum* (Pearson’s r 0.003, *p*=0.932). By considering the 24 households where people were also screened for infection, no significant association was found between having at least one person positive for any of the parasites and having a correspondent positive E/F or carcass pool (*p*<0.05).

The probability (with 95% confidence intervals) of finding a positive mosquito sample (either in the carcass or in the E/F) sampling 10 random households was 0.670 and 0.897 in Sekyerekura and Dugli respectively for *W. bancrofti*, and 0.999 for *M. perstans* in either of the two communities.

### Spatial distribution of parasite positivity in humans and mosquito samples

The spatial distribution of mosquito density, mosquito-parasite positivity, and human-parasite posivity is shown for *W. bancrofti, P. falciparum* and *M. perstans* in Figure 2, 3 and 4 respectively. No spatial clustering was observed in mosquito density in either village, and no clustering was observed in the positivity for any of the parasites in Dugli. Some clustering was observed for *W. bancrofti* in Sekyerekura, but these results should be treated cautiously as the number of households (n=24) and the limited value variation did not allow for a proper spatial analysis.

**Figure 2.**
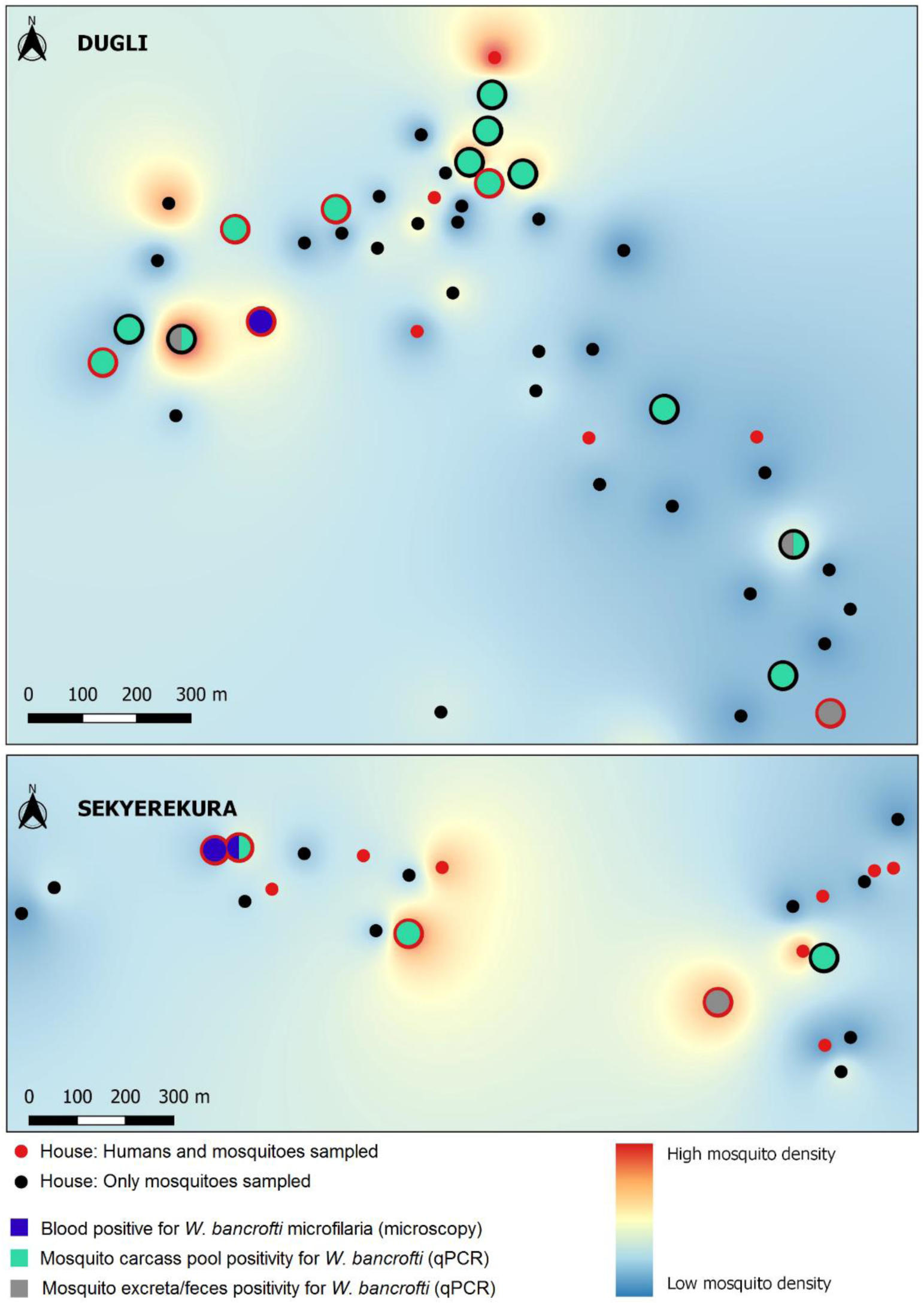
*W. bancrofti* positivity in human and mosquito samples at the household level in the communities of Dugli (top) and Sekyerekura (bottom)

**Figure 3.**
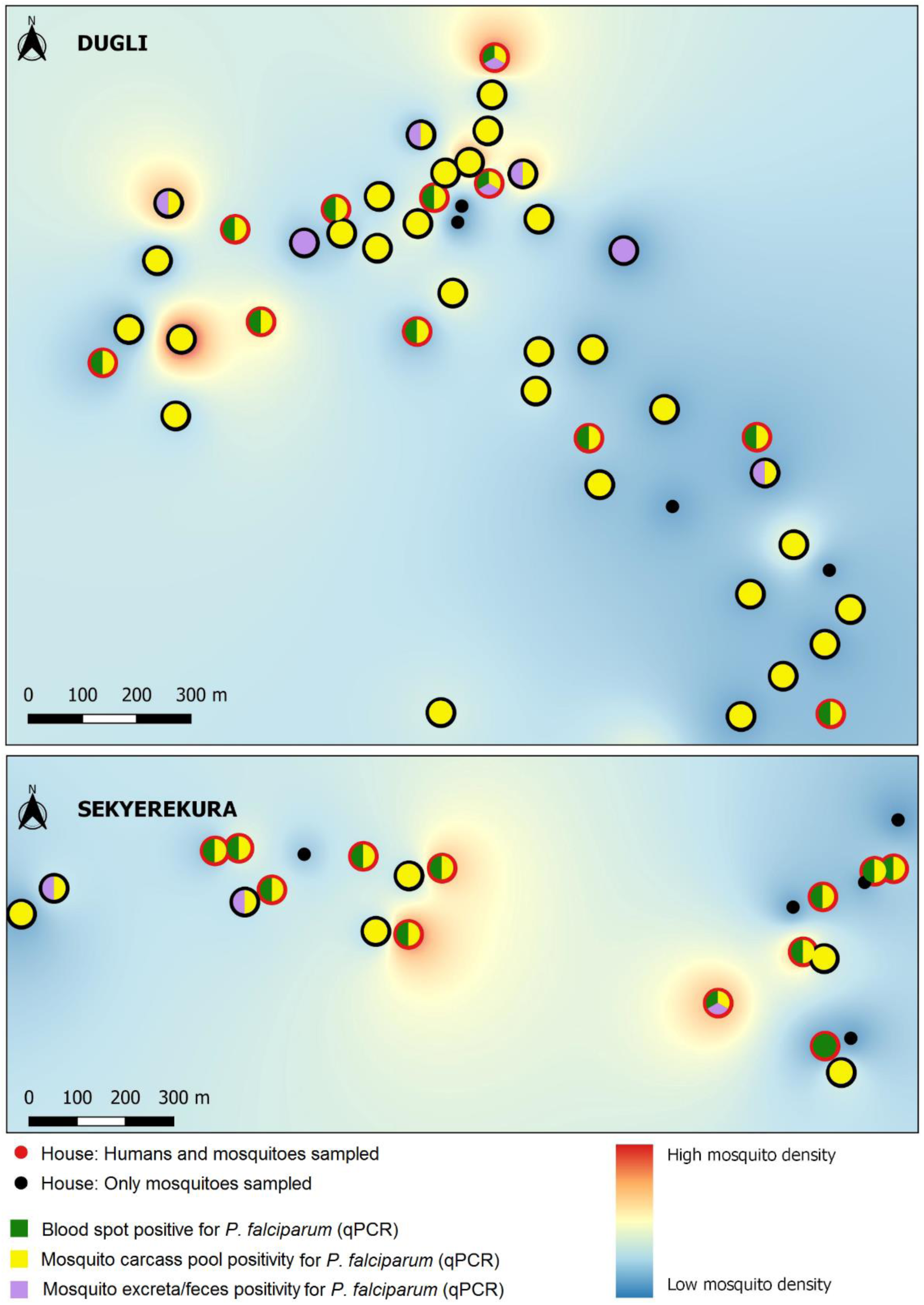
*P. falciparum* positivity in human and mosquito samples at the household level in the communities of Dugli (top) and Sekyerekura (bottom)

**Figure 4.**
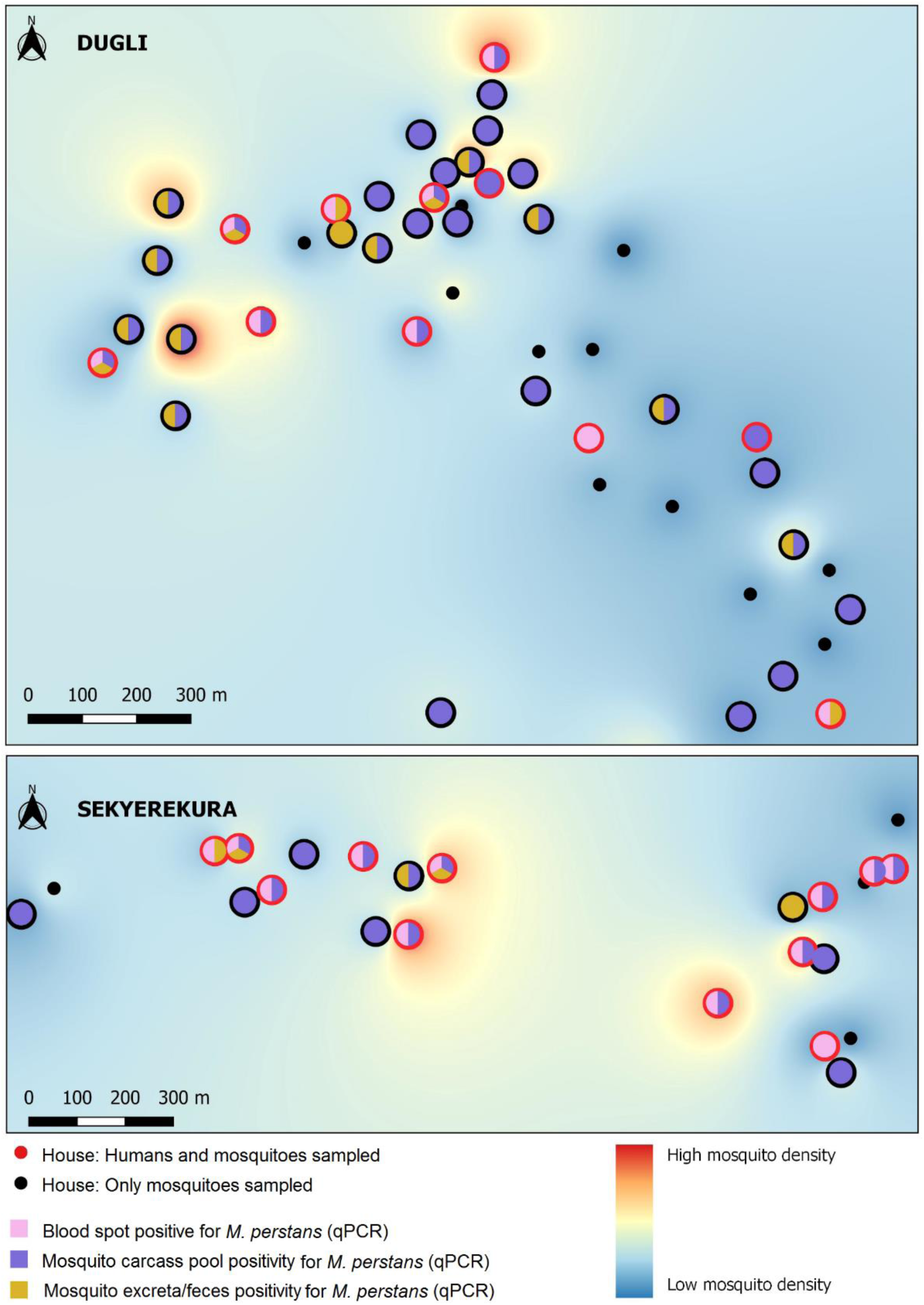
*M. perstans* positivity in human and mosquito samples at the household level in the communities of Dugli (top) and Sekyerekura (bottom)

### Evaluation of carcass cross-contamination through E/F

Experimental evaluation revealed low-levels of carcass cross-contamination as a result of the cohousing of exposed and unexposed mosquitoes (S3 Contamination results). All five experimental E/F pools allowed for *P. falciparum* detection (indicating that E/F with the potential to cross-contaminate was present and produced by the mosquitoes). However, positive results were obtained from exposed mosquitoes originating in only 2 of 5 experimental pools, with only one of the five exposed mosquitoes from each pool allowing for *P. falciparum* detection in both instances (1 H&T, 1 Abd). In conjunction, these results would suggest that nearly all parasite signal was expelled from exposed mosquitoes by the 72 hours post-exposure time point, resulting in deposition in the corresponding E/F pools. Despite this extensive deposition of potentially contaminating material, only three of five experimental mosquito pools contained an unexposed mosquito that allowed for *P. falciparum* detection upon DNA extraction. This corresponded to an overall rate of unexposed mosquito contamination of 3% (n=100).

## Discussion

In order to improve the sensitivity and scalability of MX for the detection of infection within a community, we have previously shown in the lab that mosquito E/F can be collected using a superhydrophobic insert and screened for the presence of various parasites [28]. We tested this tool in two villages of rural Ghana and we have shown for the first time that the method is fully field-applicable, including for pathogens which are not transmitted by mosquitoes.

Our primary goal was to verify the presence of *W. bancrofti* in these communities, using lymphatic filariasis as an example of the capacity of our approach to detect an infection at a very low prevalence and approaching elimination [29]. *W. bancrofti* DNA was successfully recovered from both insect E/F and carcasses, with the low DNA positivity rates comparable to the prevalence of microfilaria in people. It is important to note that the two communities underwent a round of MDA with ivermectin and albendazole two to four weeks before we initiated our study. Considering that a mosquito takes only a few microliters of blood during feeding, and that the microfilaria density estimated in our positive villagers was ranging from 0.17 and 0.04 larvae per microliter, detecting parasite DNA in mosquitoes and their E/F shortly after community drug treatment shows the high sensitivity of the approach in very low prevalence and intensity settings.

Our approach allowed for the detection of *M. perstans*, which was also the most prevalent of the pathogens in mosquito E/F. Because this filarial worm is not transmitted by mosquitoes, this finding is particularly important because it proves the feasibility of our approach to detect pathogens which are present in human blood and that are taken up with the bloodmeal by insect vectors. Our results corroborate previous findings where mosquitoes have been used to “sample” a variety of pathogens [6,8], including using the abdomens of blood-fed culicine mosquitoes to detect human malaria parasites [7]. There is scope to expand the array of pathogens circulating in the blood (including bacteria and fungi) to be tested for persistence and detection sensitivity using mosquito E/F. Interestingly, we also detected *Mansonella* DNA in the excreta of female mosquitoes collected using gravid traps.

We detected *M. perstans* DNA in the mosquito head and thorax aliquots While our experimental results show that exposure to positive E/F could result in carcass contamination, such contamination appears to occur at low levels. It is possible that the microfilaria successfully penetrated the midgut, as has been shown in laboratory infections of *Aedes aegypti* with *M. ozzardi* [44]. If *M. perstans* microfilaria manage to penetrate the midgut, the parasite may reach the thoracic muscles of the insect where the immune system would kill it and the thorax would test positive by PCR from the melanized dead larvae DNA. It is also possible that *M. persta*ns is caught in the mosquito cibarial armature during ingestion, as proposed in a previous study where DNA positivity for *B. malayi* was detected in the heads and thoraces of *Cu. pipiens* (a non-permissive vector) up to 14 days following the microfilaraemic blood meal [21]. This hypothesis seems to be corroborated by observations in our laboratory where filarial DNA is detected at 24 hours post-exposure in both the heads and the thoraces by PCR (unpublished data).

There was an unexpected and significant discrepancy in the DNA detection rate of *P. falciparum* between sample types. Despite the prevalence of malaria being very high in the communities (with at least one positive person in each sampled household), and similarly in the mosquito carcasses, the detection in the E/F was very low. One potential explanation is that our method of E/F storage (dried at room temperature without any preservative) may have caused a more rapid degradation of excreted DNA from a unicellular pathogen like *Plasmodium*, compared to whole larvae in the case of filarial worms. The possible use of collection cards for the preservation of nucleic acids, such as Whatman or GenTegra cards, as a means of overcome such potential degradation is currently being explored. Similarly, *Plasmodium* DNA could be degraded more efficiently in the mosquito midgut during blood digestion compared to filarial worms, where body remnants may be excreted in the feces still carrying enough intact DNA for the qPCR target to be detected. Another possibility is that the expulsion of *Plasmodium* DNA in the E/F of some mosquitoes did not occur during the time insects were held in the cups, effectively reducing the detection rates in the E/F compared to the carcasses.

Positivity detected in the mosquito E/F or carcasses was not predictive for the presence of infected people in the same household. In a recent study, *Plasmodium* infection incidence at the household level and the number of people infected were both correlated to the presence of *Plasmodium*-positive *Culex quinquefasciatus* collected from the same households [7]. Given the presence of *Plasmodium*-positive people in every household in our study, the predictive power of mosquito samples for malaria could not be tested. Furthermore it is not possible to predict which household members slept under a bed net the night of the collection, and whether the majority of the collected mosquitoes fed preferentially on an infected or uninfected person skewing the results. Additional and more extensive studies are needed to establish any correlation between samples, which may include repeated sampling over a longer period of time and assessing heterogeneity of exposure to bites by genotyping both the blood of the people and the blood ingested by the mosquitoes [45].

Although analysis of infection spatial clustering was performed, future research in the spatial distribution of the pathogens is needed. The limited number of positives and the low number of households (in particular in Sekyerekura) limited the tests performance. Knowledge on whether the pathogens shown an uneven spatial distribution in a community is important when planning molecular xenomonitoring sampling. Nevertheless, the probability of detection of at least a *M. perstans*-positive mosquito sample in 10 randomly selected households was higher than 95%, showing the potential for using indoor resting blood fed anophelines to detect the presence of this parasite without the need of sampling all the households.

Currently, it is clear that MX is not diagnostic but it can nevertheless aid in the verification of presence or absence of infection. Since sampling blood fed mosquitoes increases the likelihood of detection by ensuring that all sampled mosquitoes have been exposed to at least one bloodmeal, there is further scope to detect multiple bloodborne pathogens of both human and animal origin [1–3]. The possibility of detecting parasitic infections such as sleeping sickness, leishmaniasis or loiasis should be explored to assess whether MX could complement existing surveillance activities for these diseases. Our field study confirms that screening mosquito E/F and carcasses are both feasible approaches. However E/F screening can be done in larger pool sizes [22] and there are other approaches to collecting excreta, including swabbing from collection cups [28] or from passive traps [26]. Future innovations may further facilitate mosquito E/F collection in the field, for example by fitting the SHC into passive mosquito traps.

## Acknowledgements

We want to thank the communities and participants of the study who made this study possible. We also want to thank all the Ghana Health Service officers from the study districts for they invaluable support and help.

## Supporting information

**S1 Table.** Mosquito numbers and diversity in the two study communities.

| Community | Total mosquitoes | No. GA sl | % GA sl | No. FU sl | % FU sl | No. OTH An | % OTH An | No. All anophelines | % All anophelines | No. CU | % CU | No. OTH | % OTH |
| --- | --- | --- | --- | --- | --- | --- | --- | --- | --- | --- | --- | --- | --- |
| Sekyrekura | 760 | 645 | 84.9 | 79 | 10.4 | 26 | 3.4 | 750 | 98.7 | 8 | 1.1 | 2 | 0.3 |
| Dugli | 1,571 | 1,386 | 88.2 | 168 | 10.7 | 9 | 0.6 | 1,563 | 99.5 | 8 | 0.5 | 0 | 0.0 |

GA sl= *An. gambiae* sensu lato; FU sl= *An. funestus* sensu lato; OTH An= other anophelines; CU= *Culex* sp; OTH= other genera (these included one *Mansonia* sp and one which was not identified due to a damaged carcass).

**S2 Table.**
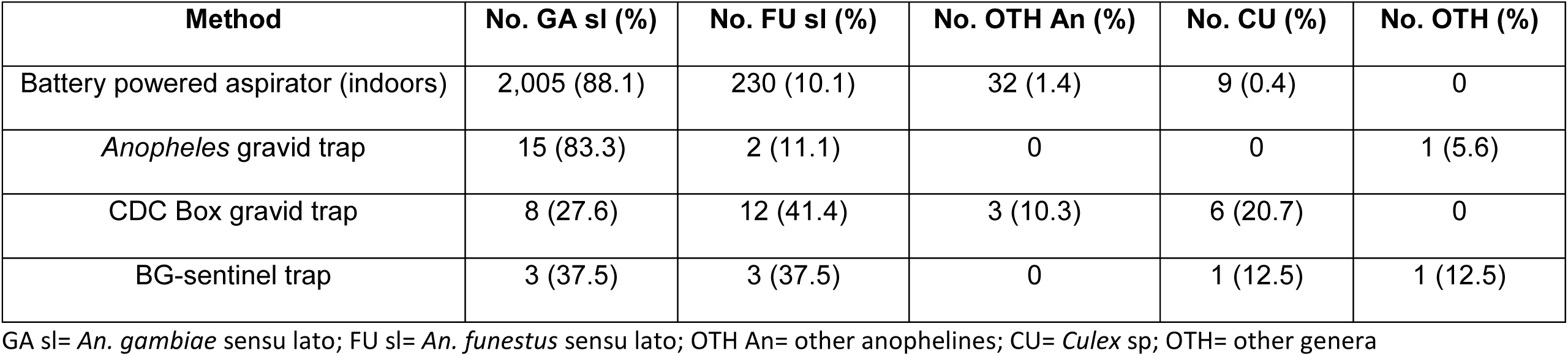
Mosquito numbers and diversity by collection method across the two communities.

| Method | No. GA sl (%) | No. FU sl (%) | No. OTH An (%) | No. CU (%) | No. OTH (%) |
| --- | --- | --- | --- | --- | --- |
| Battery powered aspirator (indoors) | 2,005 (88.1) | 230 (10.1) | 32 (1.4) | 9 (0.4) | 0 |
| *Anopheles* gravid trap | 15 (83.3) | 2 (11.1) | 0 | 0 | 1 (5.6) |
| CDC Box gravid trap | 8 (27.6) | 12 (41.4) | 3 (10.3) | 6 (20.7) | 0 |
| BG-sentinel trap | 3 (37.5) | 3 (37.5) | 0 | 1 (12.5) | 1 (12.5) |

GA sl= *An. gambiae* sensu lato; FU sl= *An. funestus* sensu lato; OTH An= other anophelines; CU= *Culex* sp; OTH= other genera

**S3 Supporting Information.**
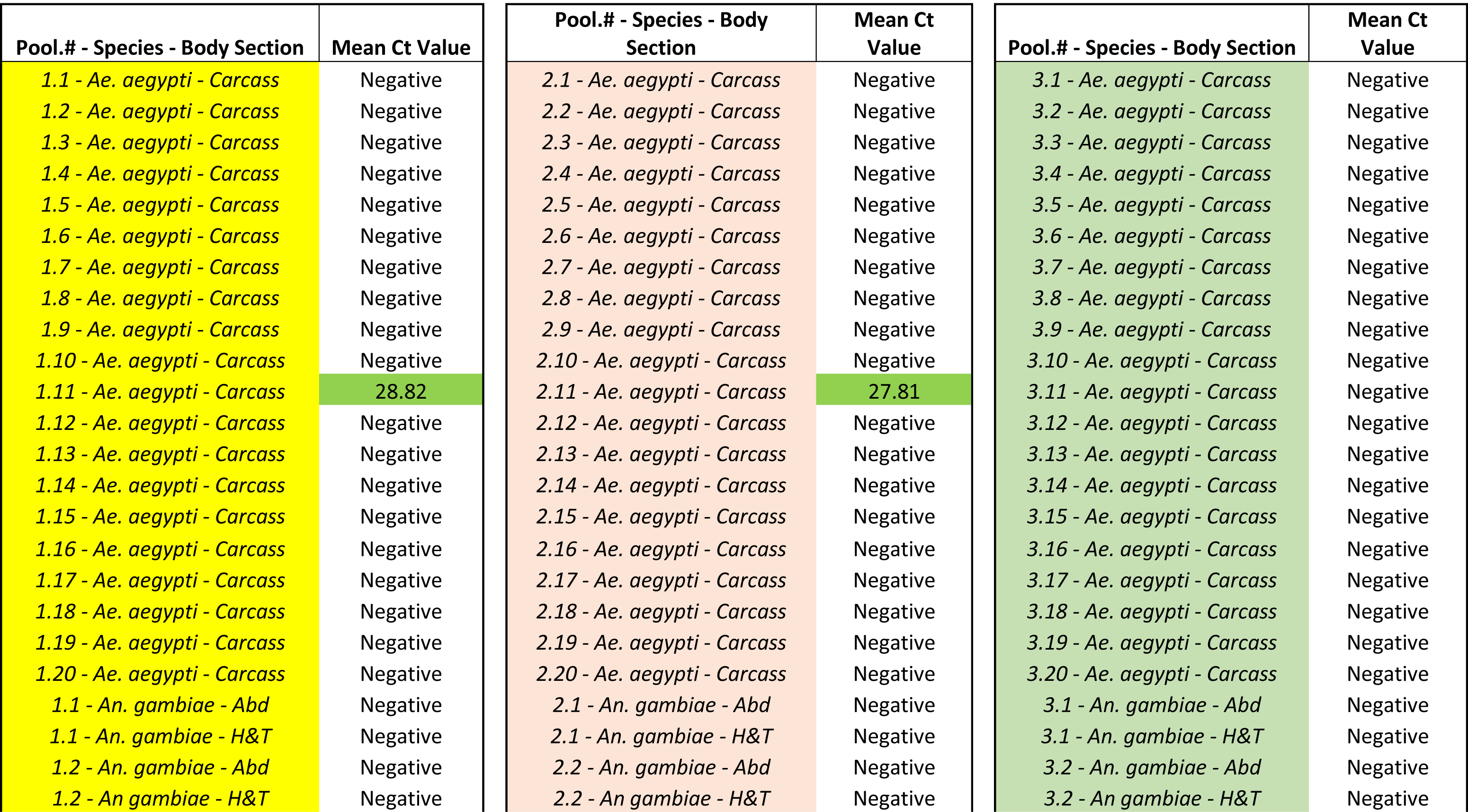
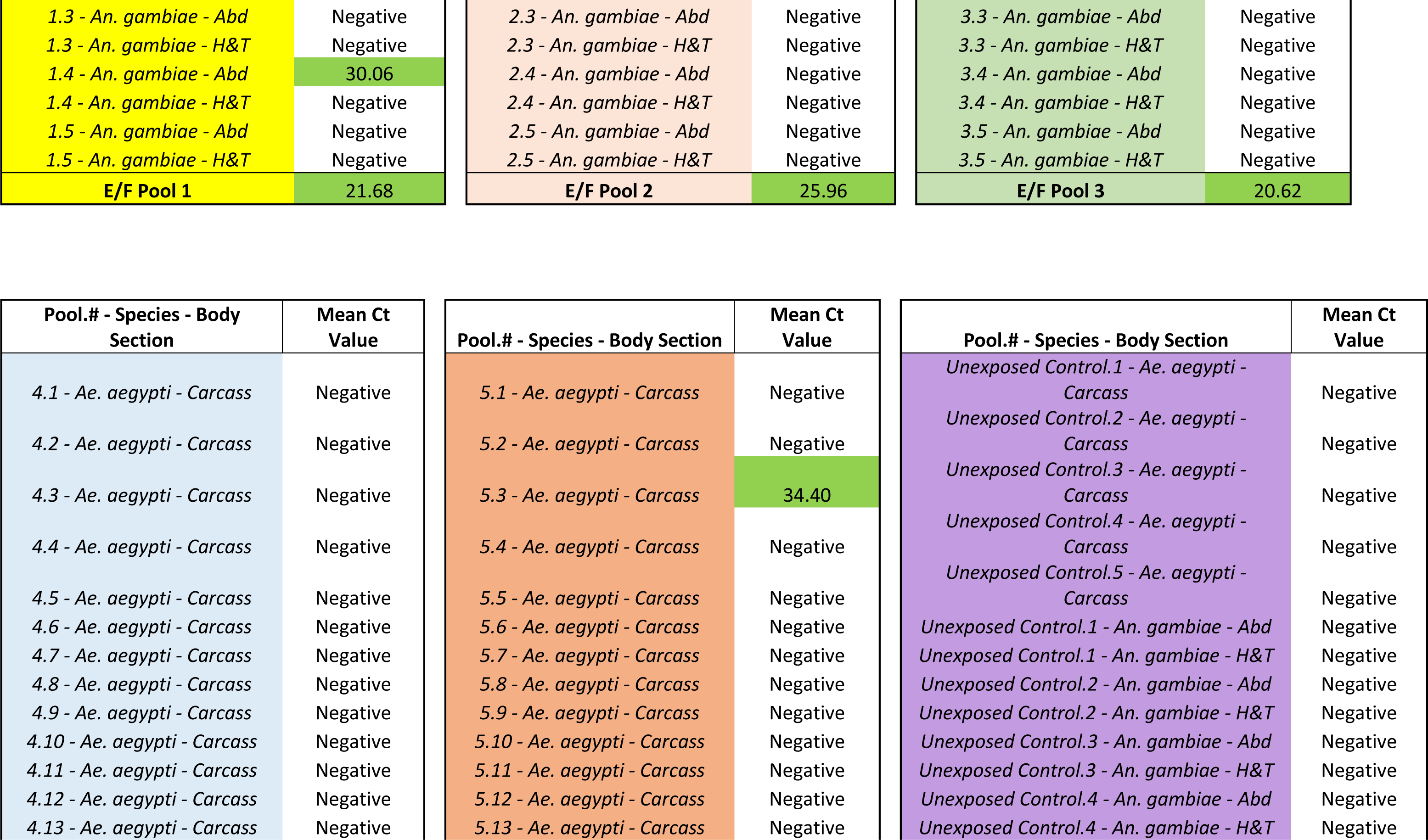
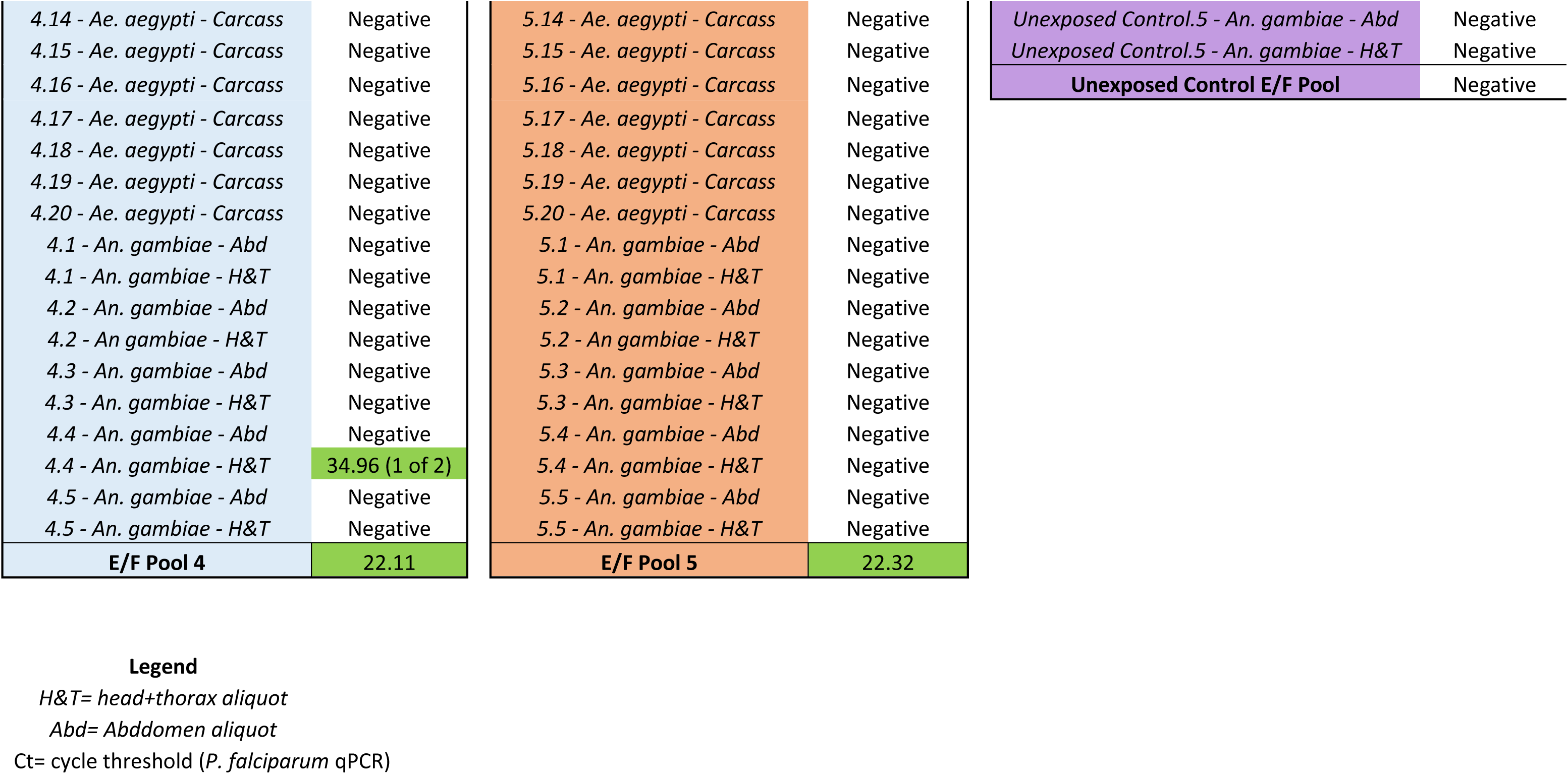
Contamination experiments results.

